# A User-Friendly Ear-EEG-Based Brain-Computer Interface Using Text Sequence Stimulation

**DOI:** 10.64898/2026.05.15.721815

**Authors:** Xiaoyang Li, Zhuoran Xu, Bowen Li, Yijun Wang, Xiaorong Gao

## Abstract

**Background:** Ear-EEG-based brain-computer interfaces (BCIs) provide improved wearability and comfort compared to traditional scalp-EEG systems. However, their performance is constrained by low signal-to-noise ratios (SNRs) and high rates of BCI illiteracy under conventional luminance-modulated steady-state visual evoked potential (SSVEP) paradigms.

**Methods:** This study introduces a text-sequence stimulation paradigm to address these limitations by leveraging ventral visual pathway responses that are more accessible to electrodes near the ear. Using offline frequency-sweeping experiments across 4–8 Hz, we identified optimal stimulus parameters (4.6–6.8 Hz with 0.25π phase shifts) and integrated them into a 12-target BCI system. We further conducted online experiments to compare the response characteristics and real-time spelling performance between the proposed text-sequence paradigm and conventional luminance stimulation.

**Results:** Comparative experiments with 14 participants demonstrate that text sequence stimuli achieve an average information transfer rate (ITR) of 44.59 ± 10.50 bits/min, outperforming luminance modulation by 76.18% in ITR. Notably, text sequence stimulation effectively mitigated BCI illiteracy, with all participants achieving near or above 70% accuracy (mean: 86.37 ± 9.61%). This represents a significant improvement over luminance modulation, where 50% of users fell below 70% accuracy.

**Conclusions:** By reducing the flicker area by 14% and mimicking the natural luminance variations that occur during reading, the proposed method enhanced visual comfort. The online results further validate text-sequence stimulation as a high-performance and user-friendly paradigm for ear-EEG BCIs, supporting their practicality for assistive applications.

## 1 Introduction

Brain–computer interfaces (BCIs) convert neural activity into computer commands in real time, opening alternative communication pathways for users[1,2]. Non-invasive electroencephalography (EEG) remains the workhorse of BCIs because of its portability and millisecond temporal resolution[3]. Beyond clinical and laboratory use, a major goal is to enable consumer-grade, everyday BCI applications (e.g., wearable human–computer interaction) that are fast to set up and socially acceptable. Yet conventional scalp-EEG systems still require gel-based electrode caps or headbands, causing lengthy setup, aesthetic concerns, and user discomfort[4–6].

Ear-EEG, which records from hair-free regions around the auricle, has emerged as a practical alternative, enabling quick mounting suitable for everyday tasks such as visual or auditory decoding and fatigue monitoring[4,6–8]. With ear-electrode BCIs, steady-state visual evoked potentials (SSVEPs) are especially attractive for multi-target BCIs (e.g., virtual keyboard spellers) because they require little user training and can achieve high information transfer rates (ITRs)[4,7]. At present, ear-electrode SSVEP BCIs can reach a maximum ITR of 37.5 bits/min[4].

However, luminance modulated SSVEPs suffer from two major drawbacks that are critical in ear-EEG applications. First, the neural response is highly dependent on stimulus intensity, prolonged fixation on such luminance stimulation inevitably leads to visual fatigue[9–11]. Second, luminance flicker predominantly activates neural sources in the occipital lobe, resulting in a spatial mismatch with ear electrodes[12– 15]. These limitations jointly reduce the signal-to-noise ratio (SNR) and keep the ITR far below scalp-EEG benchmarks. Moreover, such constraints are associated with the “BCI illiteracy” phenomenon observed in a subset of users[12,16]. In this work, BCI illiteracy is operationally defined as a classification accuracy below 70%[17].

Recent work has explored alternative stimulation strategies. Color modulated stimuli recruit ventral pathway regions to boost mid-and high-frequency responses, enhancing comfort[18]. Depth-frequency modulation has achieved 91 % accuracy in two-target systems[19]. And bilateral phase-coding improves SNR by applying distinct phase shifts to each hemifield[12]. Yet such paradigms often trade target capacity or ITR for ergonomic gains and still leave the core spatial-alignment problem unsolved.

Text sequences represent a promising alternative for improving ear-EEG based SSVEP BCIs. During visual text recognition, a robust event-related component around 170 ms (N170) is consistently elicited and is closely related to orthographic processing along the ventral visual pathway[20], [21], [22]. This differs fundamentally from conventional luminance-modulated SSVEP, which relies primarily on global luminance oscillations and predominantly drives occipital responses. Importantly for ear-EEG recordings, such activity exhibits a spatial distribution that can be more accessible to electrodes placed around the auricle/mastoid region than purely occipital responses. This pathway not only lies in closer spatial proximity to ear-EEG electrodes but is also relatively insensitive to variations in stimulus contrast and size, allowing stable responses to be elicited under naturalistic viewing conditions[23,24]. Moreover, text stimuli require lower luminance intensity, reducing visual fatigue and enhancing overall user comfort. Taken together, these properties suggest that text-evoked SSVEPs can effectively raise SNR, alleviate fatigue, and substantially improve the usability and performance of ear-EEG BCIs in practical applications.

In this study we introduce a text-sequence stimulation paradigm designed to harmonize neural activation with ear-EEG’s anatomical constraints. We define text-sequence stimulation as a joint frequency–phase coded visual stimulation paradigm in which each target presents a rapid sequence of different text. The text updates are synchronized to the assigned frequency and phase, so that attending to a target produces frequency-tagged responses (fundamental and harmonics) while repeatedly engaging orthographic processing mechanisms. First, we characterized the response properties of text-sequence stimulation in offline experiments, and optimized the stimulus parameters and decoding pipeline accordingly. Subsequent online experiment compared text and luminance paradigms in a 12-target spelling task. With 3 s stimulation, the text paradigm achieved 44.59 ± 10.50 bits/min at 86.37 ± 9.61 % accuracy, outperforming conventional luminance modulation by 76.18 %. Remarkably, every participant accuracy near or exceeded 70 % with text stimuli, eliminating the “BCI-illiteracy” issue common in Ear-EEG BCIs. These results underscore the potential of text stimulation for practical, high-ITR ear-EEG BCIs.

In particular, such improvements are highly relevant for assistive communication and rehabilitation-oriented applications, where rapid setup, prolonged wearing comfort, and stable performance are critical. Compared with conventional scalp-EEG systems, ear-EEG offers practical advantages for bedside communication and caregiver-assisted use in populations with severe motor impairments, such as amyotrophic lateral sclerosis or brainstem stroke. In this context, the present 12-target spelling task is intended as a proof-of-concept rather than a fully optimized language interface, demonstrating the feasibility of paradigm-level stimulus design for improving the performance–usability trade-off of ear-EEG BCIs.

## 2 Methods

### Participants

Fifteen healthy adults (7 female; mean age: 27 years, range: 20-35) participated in the offline frequency-sweeping experiment. Fourteen participants (7 female; mean age: 28 years, range: 20-57) subsequently completed the 12-target online BCI spelling task, with7 participants overlappingbetween the two experiments. To better evaluate the system’s adaptability, we did not impose restrictions on age, sex, or prior BCI/EEG experience during online experiment recruitment. One participant (female, 57 years) represented an outlier in age, while others fell within a narrower range (20-33 years). To mitigate order effects, participants were divided into two groups for the online experiment: half performed the luminance modulated task first, while the other half began with the text sequence stimulation paradigm. All participants were native Chinese speakers with normal or corrected-to-normal vision. The study was approved by the Tsinghua University Ethics Committee, and informed consent was obtained.

### Data Recording

Experiments were conducted in a well-lit room, with participants seated 70 cm from a monitor. EEG data were recorded using Synamp2 system at 1000 Hz. Full scalp EEG (62 channels) following the 10-20 system and ear-EEG (8 channels) were collected, electrodes location as shown in Fig. 1A, actual ear-electrode placement as shown in Fig. 1B.

**Fig. 1.**
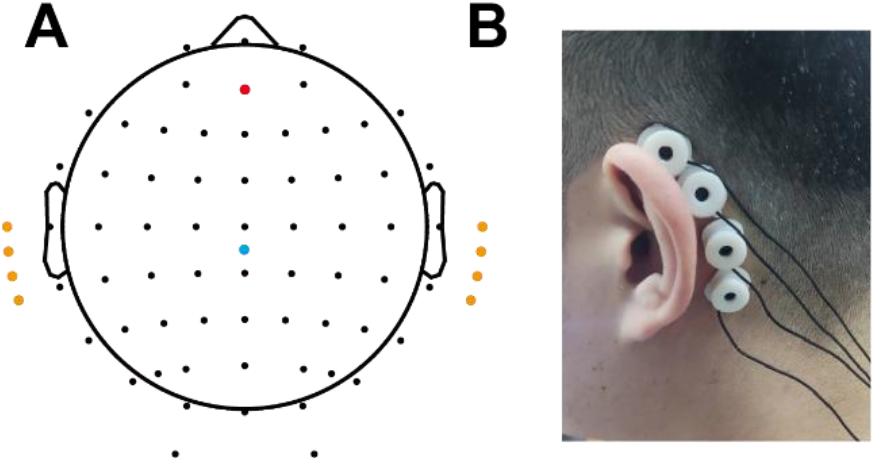
Electrode montage. (A)Black dots indicate the 62 scalp electrodes. The blue dot marks the reference electrode placed at the vertex, and the red dot marks the reference electrode placed at AFz. The yellow dots represent the eight ear-EEG channels, extending from the upper auricle down to the region near the mastoid. (B) Actual ear-electrode placement.

### Offline Frequency-Sweeping Experiment

To facilitate the development of a text-based ear-EEG BCI system, we conducted a single-target frequency-sweeping experiment to investigate EEG responses to text stimuli from 4 to 8 Hz with a 0.2 Hz resolution. Each trial consisted of a 1 s fixation cross, 8 s text stimulation, and a 1 s rest period, as shown in Fig. 2A. Text stimulation was implemented as a sequential presentation of Chinese characters, where glyphs switched at predefined frequencies (4.0, 4.2, …, 8.0 Hz). Characters were updated at the rising edges of the waveform with a phase of 0, as defined in Eq. (1). The text stimuli consisted of 100 commonly used Chinese characters, and the stimulus size did not exceed 140 × 160 pixels (width × height). Each character appeared with equal probability in a randomized order. The experiment comprised 12 blocks, each containing 21 trials corresponding to the 21 distinct stimulation frequencies. Within each block, the order of stimulation frequencies was randomized, while the character sequence for a given stimulation frequency was kept identical across repetitions.

**Fig. 2.**
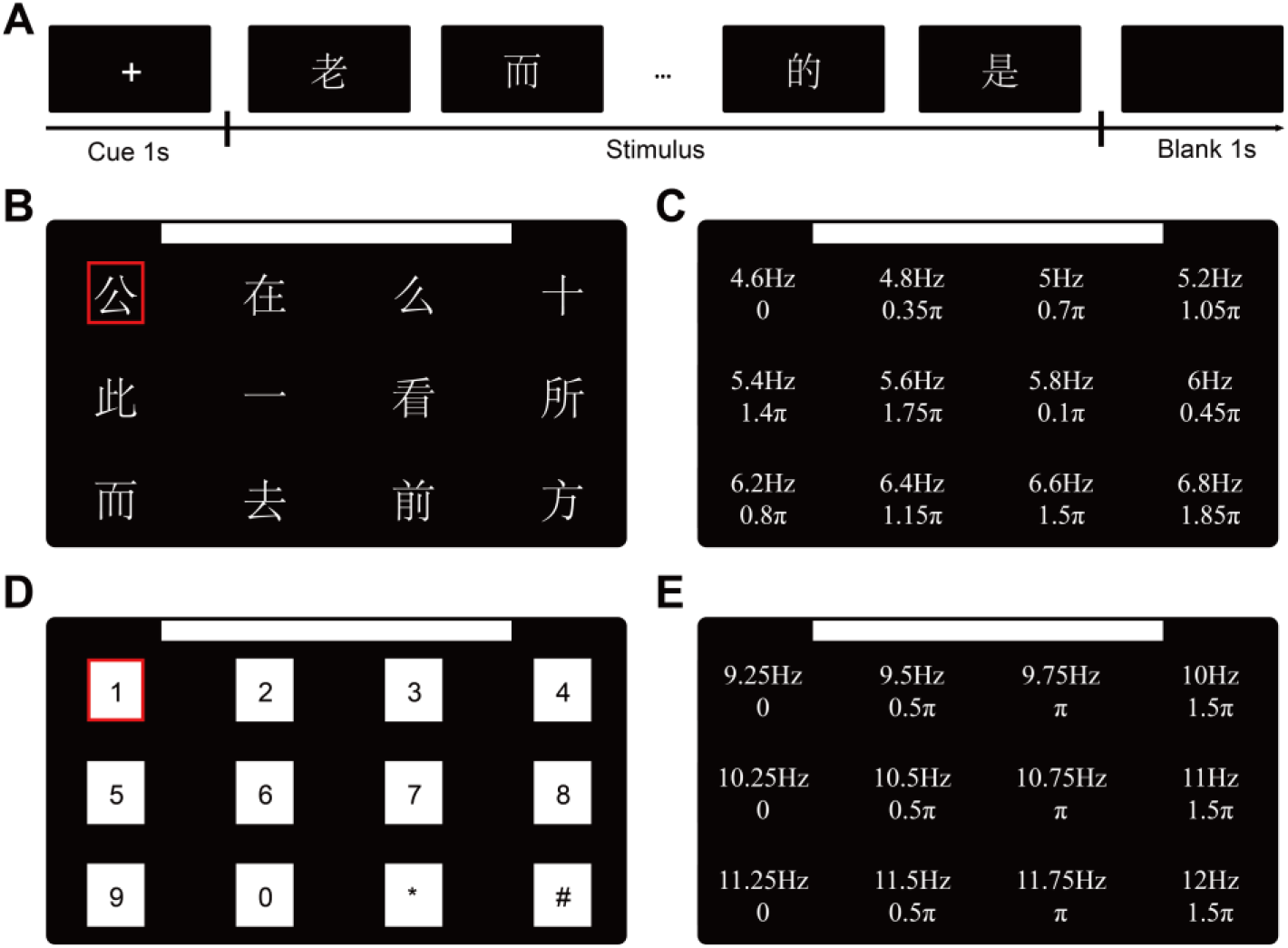
Stimulation paradigm. (A) schematic of a single trial stimulus process in the offline experiment. (B) 12 targets interface for text sequence stimulation in online experiments. The boundary of the stimulus represented by the red box. (C) Frequency phase encoding of text stimuli. (D) 12 targets interface for luminance stimulation in online experiments. The same boundary of the stimulus represented by the red box. (E) Frequency phase encoding of luminance stimuli.

Stimuli were programmed usingMATLAB 2015band thePsychtoolbox, ensuring precise temporal synchronization. Visual stimuli were displayed on a 24.5-inch LCD monitor with a resolution of1920 × 1080 pixelsand a refresh rate of240 Hz, critical for maintaining phase-locked flicker accuracy.

### Online Experiment

We conducted an online experiment to compare the BCI performance of the proposed text-sequence paradigm with a validated luminance-modulated paradigm whose parameters were optimized in prior ear-EEG studies. Both paradigms used the frequency–phase encoding scheme illustrated in Fig. 2B–E. For luminance modulation, the parameters followed those established in previous ear-EEG BCI studies: frequencies ranging from 9.25 to 12 Hz (with 0.25 Hz intervals) and a phase interval of 0.5π. This frequency-phase encoding has been shown in published literature to yield the highest ITR[4]. The text paradigm utilized offline-optimized parameters: frequencies 4.6–6.8 Hz (0.2 Hz intervals) and phase interval of 0.25π. All targets were simultaneously modulated with unique frequency-phase combinations during stimulation. Text and luminance stimuli sizes were matched, with text characters confined within the borders of luminance stimuli (red in Fig. 2B and Fig. 2D), with a bounding box of 140 × 160 pixels (width × height), consistent with the offline experiment.

In text sequence stimulation, visual flicker followed a square-wave modulation (Fig. 2C), where characters updated at rising edges of the waveform:

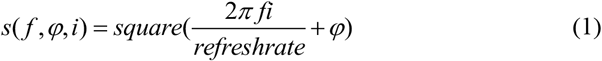

In luminance modulation, luminance varied sinusoidal waveform (Fig. 2E):

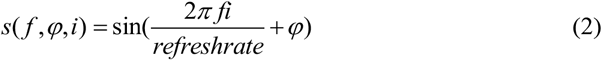

where *f* represents stimulus frequency, *ϕ* represents initial phase, and *i* represents frame index.

To mitigate potential order effects (e.g., learning or fatigue), participants were counterbalanced across paradigms for two groups: seven participants completed the text-sequence paradigm first, while the remaining seven completed the luminance paradigm first. Each participant completed 12 training blocks to train the classification model, followed by 10 testing blocks to evaluate online performance. Each block consisted of 12 trials, covering all 12 targets once per block with a randomized target order. During the training phase (12 blocks), each trial comprised a 0.5 s cue indicating the target to attend, a 5 s visual stimulation period, and a 0.5 s dummy feedback interval with the next cue. During the testing phase (10 blocks), each trial comprised a 0.5 s cue, a 3 s visual stimulation period, and a 0.5 s feedback period displaying the recognized output in the interface with the next cue.

The software/hardware implementation and stimulation setup in the online experiment were kept consistent with those used in the offline experiments. Across the two paradigms, hardware configuration, interface layout, trial timing, number of targets, training and testing protocol, and decoding framework were kept identical. Differences in stimulation frequency bands, phase intervals, and waveform shapes were specific to each paradigm, reflecting their distinct physical implementations and dominant neural response mechanisms. As a result, paradigm-matched parameter settings were adopted to characterize the achievable performance under each stimulation strategy.

### Data Evaluation

To account for inherent visual system latency, EEG data epochs were extracted from 0.14 s to 5.14 s post-stimulus, with the initial 140 ms offset compensating for visual processing delays[14]. The extracted data were downsampled to 250 Hz and notch-filtered to remove powerline interference. A 4th-order Butterworth bandpass filter (2–90 Hz) was applied to isolate frequency components of interest.

Narrowband SNR was computed as the ratio of the amplitude at the target frequency to the mean amplitude of adjacent frequency bins:

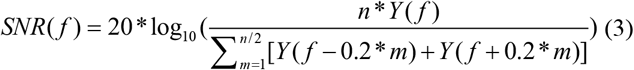

where *f* is the target frequency,*Y(·)* represents amplitude spectrum, and *n*=10 (i.e., ±1 Hz around *f*). For ear-EEG, SNR was averaged across 8 posterior auricular channels; for scalp EEG, 21 channels were used (Pz, P1, P2, P3, P4, P5, P6, P7, P8, POz, PO3, PO4, PO5, PO6, PO7, PO8, Cb1, Oz, O1, O2, Cb2).

### Decoding Algorithm and Validation

The task-discriminant component analysis (TDCA) algorithm was employed due to its superior performance[25]. TDCA integrates data augmentation, discriminant spatial pattern extraction, and sub-spatial filter optimization to enhance signal separability. Foroffline validation, leave-one-out cross-validation was applied to evaluate 12-class classification performance. Inonline testing, 12 blocks served as training data, while 10 blocks were used for real-time performance assessment. For the luminance-modulated condition, we used the same decoding method and parameter settings as the highest-ITR system reported in prior work[4]. For the text-sequence condition, we employed a filter-bank configuration optimized specifically for text stimulation. All other software/hardware settings and experimental parameters were kept consistent across the two paradigms to enable a fair comparison of performance between paradigms. Decoding was performed using all ear-EEG channels.

### Filter Banks

Filter banks are widely utilized in SSVEP decoding due to the pronounced amplitude differences of EEG signals across harmonic frequency components. By decomposing the signal into sub-bands, filter banks enhance the classification utility of high-frequency components. Previous studies predominantly adopted theM2 filter bank design, which partitions the frequency spectrum into predefined sub-bands[25–28].

However, SSVEP responses in ear-EEG can exhibit spectral characteristics different from scalp EEG. To account for this, we implemented two standard filter-bank configurations for sub-band decomposition (M1 and M2), as illustrated in Fig. 3. Let *frequency*_*low*_ and *frequency*_*high*_ denotes the minimum and maximum stimulation frequencies in the encoding set, *K* denotes the number of sub-bands, and *k*∈{1,…,*K*} denotes the sub-band index. In M1, the *k*-th sub-band passband is defined as

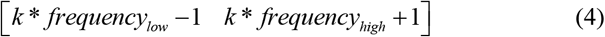

where the ±1 Hz provides a margin to reduce sensitivity to frequency leakage. In M2, the *k*-th passband is defined as:

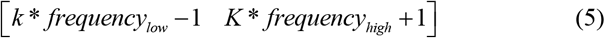

**Fig. 3.**
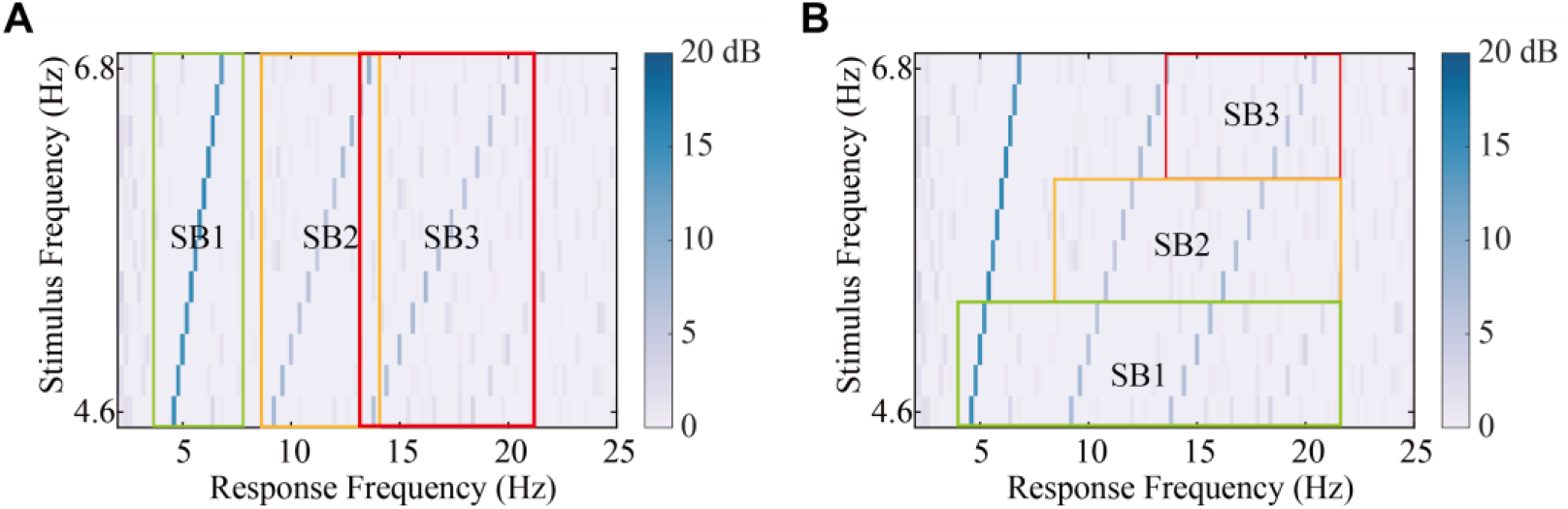
Filter banks configuration illustrated using the offline text frequency-sweeping results. The y-axis (Stimulus Frequency) lists stimulation frequencies parameters used in the online experiment (4.6 –6.8 Hz), where each row corresponds to one stimulation frequency. The x-axis (Response Frequency) indicates candidate response-frequency components (fundamental and harmonics). Colors show ear-EEG SNR at each response frequency. SB1–SB3 denote the sub-band regions covered by the corresponding filter banks in (A) M1 and (B) M2.

so that all sub-bands share the same upper cutoff to cover response components up to the highest modeled harmonic. For the text paradigm (*frequency*_*low*_=4.6, *frequency*_*high*_=6.8) the first sub-band is [3.6, 7.8] in M1 and [3.6, 21.4] in M2.

Sub-band signals were processed using TDCA spatial filters, and weighted correlation coefficients derived from these sub-bands served as features for target identification:

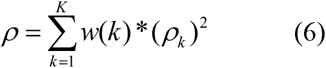

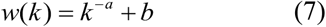

where*ρ*_*n*_ represents the correlation coefficient of the *n*-th sub-band, and *w(n)* denotes its corresponding weight. In the optimized configuration, the sub-band weighting coefficients were set to a = 0.2 and b = 0.3, yielding improved classification accuracy over other settings.

### Performance Evaluation

The decoding performance of BCI system was evaluated using two metrics: classification accuracy and ITR. The ITR was calculated according to Equation (8):

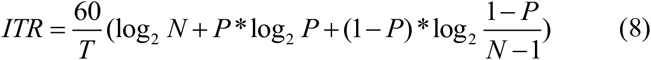

where *N* represents number of targets in the BCI system, *P* represents online classification accuracy, *T r*epresents total recognition time, defined as the data duration used for classification plus a 0.5-second gaze-shifting interval. In this study, the system was configured with *N*=12targets. The recognition time *T* incorporated both the 3s stimulus-specific data window and the 0.5s fixed gaze-shifting period, ensuring realistic operational conditions.

### Questionnaire experience evaluation

We conducted a subjective evaluation of the proposed text sequence stimulation paradigm (Fig. 2B) by comparing it to a luminance paradigm (Fig. 2D). A questionnaire was administered to assess preference level, comfort level, and perception of stimulus flicker, each rated on a 5-point scale (1–5). The preference level was rated as follows: (1)very disgusting, (2) disgusting, (3) neutral, (4) likable, (5) very likable. The comfort level was rated as: (1) very uncomfortable, (2) uncomfortable, (3) slightly uncomfortable, (4) comfortable, (5) very comfortable. The flicker level was rated as: (1) very annoying, (2) annoying, (3) slightly annoying, (4) perceptible, (5) imperceptible. Lower scores reflected stronger flicker perception, lower comfort, and lower preference[9,29–31]. All subjects who participated in the online experiment completed the questionnaire.

## 3 Results

### Response Characteristics

The offline results are shown in Fig. 4. Fig. 4A presents the SNR spectrum of the EEG (scalp) and ear-EEG responses under text stimulation at 6 Hz. While clear peaks are observed at the fundamental and lower-order harmonics, the ear-EEG SNR becomes much weaker at higher harmonics, and is nearly negligible beyond the third harmonic. As shown in Fig. 4B, when sweeping the stimulation frequency from 4 to 8 Hz, the ear-EEG SNR of the fundamental component first increases and then decreases. In contrast, the SNR of the second harmonic shows an opposite trend, decreasing at lower frequencies and increasing at higher frequencies.

**Fig. 4.**
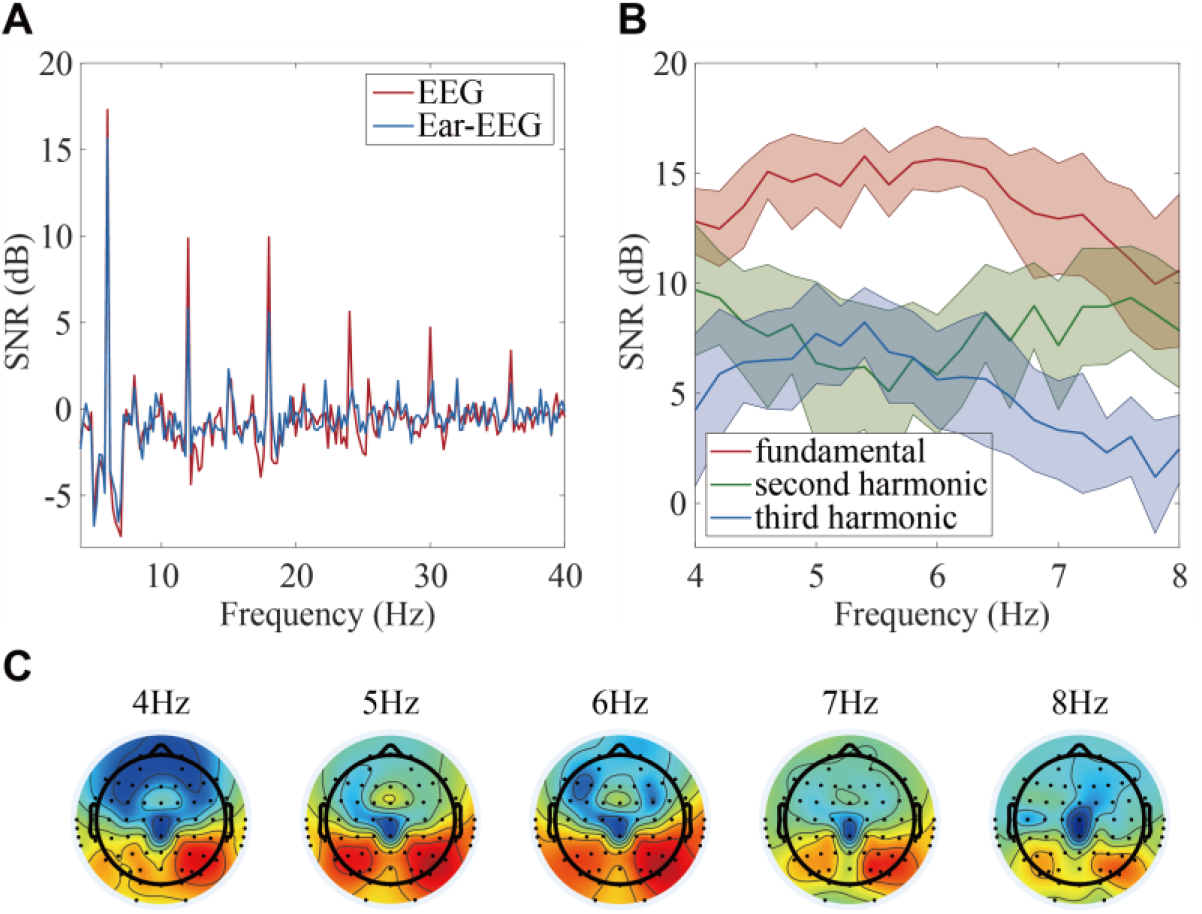
Response characteristics of text stimulation in offline experiment. (A) Frequency spectrum of 6Hz text stimulation. (B) Variation of ear-EEG SNR at the fundamental, second harmonic, and third harmonic as a function of stimulation frequency. (C) Topographic maps of responses elicited by text stimulation from 4 to 8 Hz.

Fig. 4C further illustrates the spatial distribution of the text-evoked response across frequencies. A more pronounced ventral-stream–like pattern is observed around 5– 6 Hz, which becomes less prominent as the stimulation frequency increases. Notably, this favorable range is consistent with the temporal scale of the N170 component in visual text processing: 1000/170≈5.88 Hz, suggesting that stimulation near ∼6 Hz may better engage N170-related processing and yield stronger responses accessible to ear-area electrodes.

The response characteristics of text-sequence and luminance stimulation are summarized in Fig. 5 and Fig. 6. FFT-based analysis showed that both paradigms elicited clear fundamental and harmonic components in ear-EEG, confirming that reliable steady-state responses can be generated under both stimulation types. Across all harmonics, SNR values were consistently higher in scalp EEG than in ear-EEG. Because the two paradigms used different optimized encoding bands in this 12-target speller, their harmonic spacing and potential spectral overlap differed accordingly. Specifically, the luminance band (9.25–12 Hz) yields relatively larger harmonic separation within the analyzed range, whereas in the text band (4.6–6.8 Hz) the higher-order components become closer and partial overlap can emerge from the second harmonic onward.

**Fig. 5.**
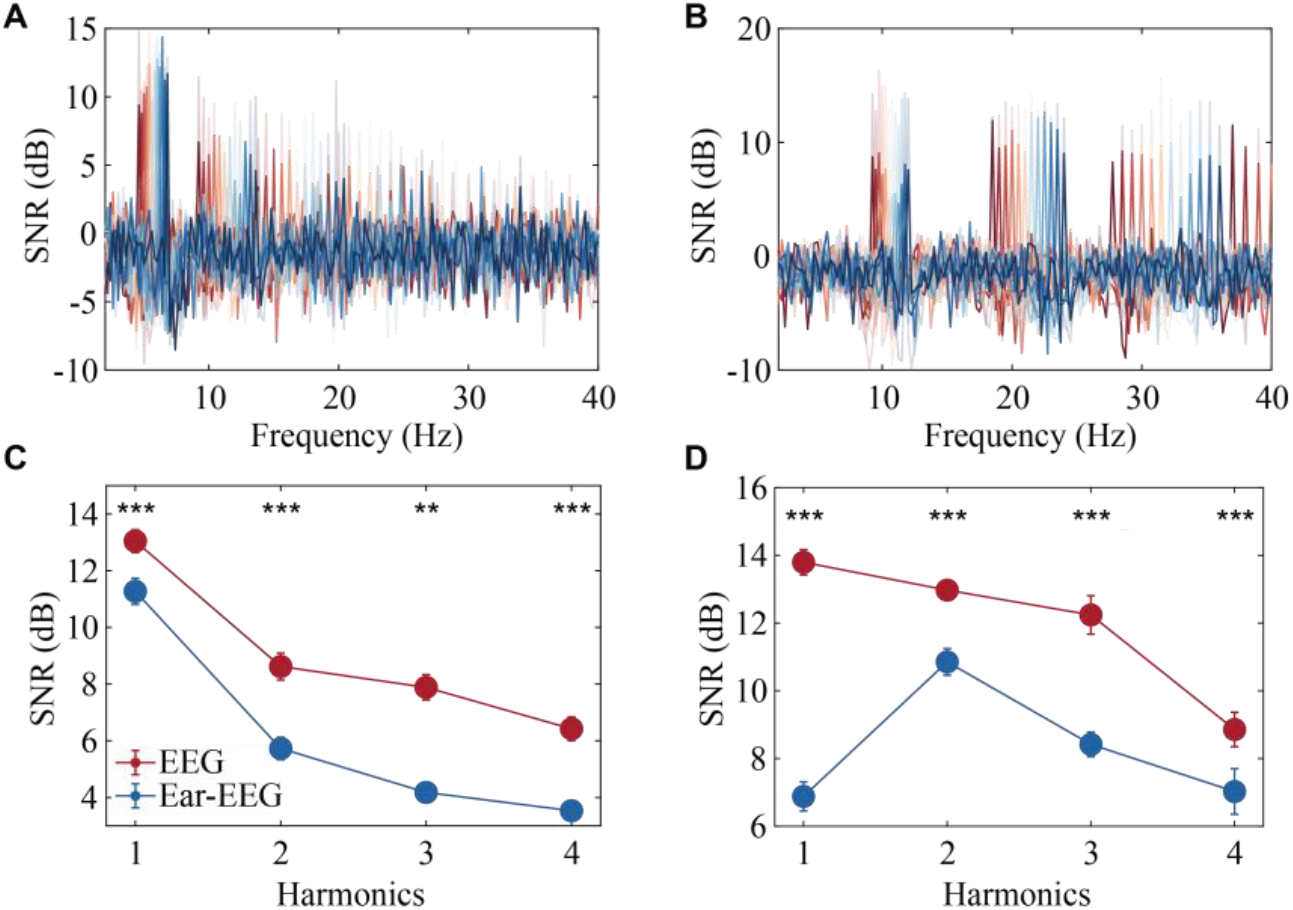
Response characteristics of text and luminance stimulation in online experment. (A) Frequency spectrum of text-evoked SSVEP responses (4.6–6.8 Hz). (B) Frequency spectrum of luminance-evoked SSVEP responses (9.25-12 Hz), different colors indicate different stimulation frequencies. (C) Harmonic SNR comparison between scalp EEG and ear-EEG under text stimulation. (D) Harmonic SNR comparison between scalp EEG and ear-EEG under luminance stimulation. Error bar represents the standard error between frequencies (paired t-test between EEG and ear-EEG, **p<0.01, *** p<0.001).

**Fig. 6.**
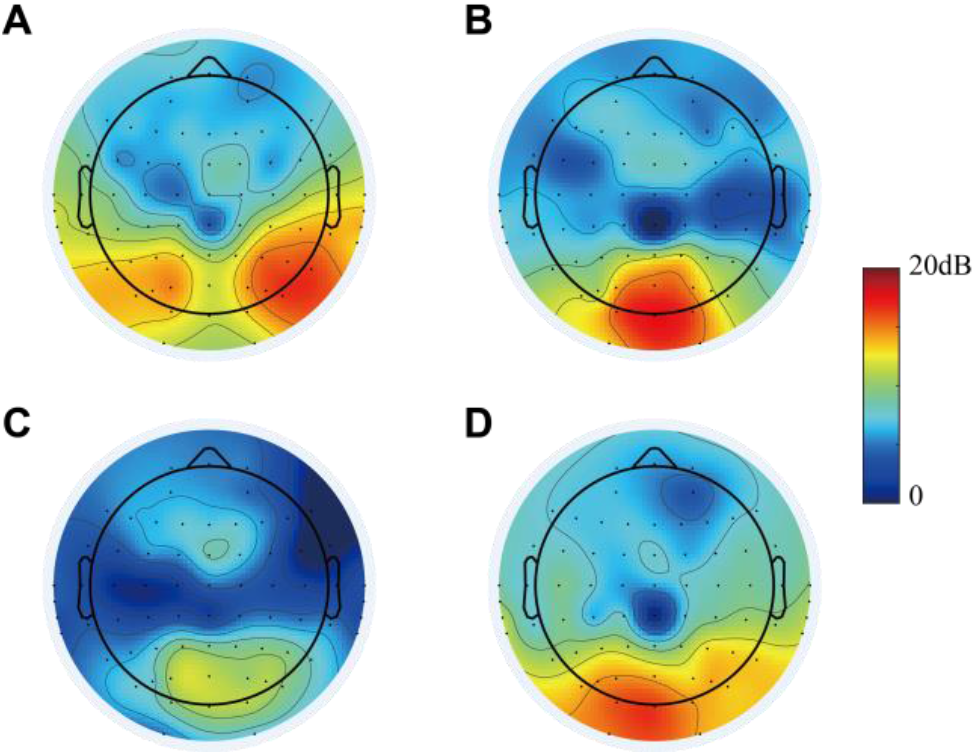
Response SNR topographic of text and luminance stimulation in online experment. (A)Topographic distribution of fundamental responses at 5.4 Hz under text stimulation. (B) Topographic distribution of fundamental responses at 11 Hz under luminance stimulation. (C) Topographic distribution of second harmonic responses at 5.4 Hz under text stimulation. (D) Topographic distribution of second harmonic responses at 11 Hz under luminance stimulation.

Within the 4.6–6.8 Hz encoding range of the text paradigm, ear-EEG exhibited robust SSVEP responses: at the fundamental frequency, the ear-EEG SNR was only 1.77 dB lower than scalp EEG (see the corresponding panel), supporting the feasibility of ear-EEG for steady-state decoding in this low-frequency band. As harmonic order increased, SNR decreased in both scalp and ear-EEG, with scalp–ear differences of 2.89 dB, 3.71 dB, and 2.89 dB at the second, third, and fourth harmonics, respectively.

Notably, the two paradigms showed different harmonic trajectories. Under luminance stimulation, scalp EEG exhibited a monotonic decrease in SNR with harmonic order, whereas ear-EEG showed an initial rise followed by a decline, with the peak occurring at the second harmonic. At the fundamental frequency, luminance stimulation yielded the largest scalp–ear discrepancy (6.91 dB), which decreased to 2.12 dB at the second harmonic; the corresponding differences at the third and fourth harmonics were 3.82 dB and 1.83 dB, respectively. These distinct patterns indicate that the harmonic structure observed in ear-EEG is not a simple attenuated version of scalp EEG and may reflect differences in the dominant neural contributions captured by the two electrode types.

We further compared SNR between paradigms. For scalp EEG, the text paradigm showed a small, non-significant difference at the fundamental frequency (−0.75 dB, p > 0.05) but significantly lower SNR than luminance stimulation at the 2nd–4th harmonics, with differences of −4.35 dB, −4.35 dB, and −2.41 dB (all p < 0.001). For ear-EEG, text-sequence stimulation produced a significantly higher fundamental SNR than luminance stimulation (+4.39 dB, p < 0.001), whereas at the 2nd–4th harmonics it was significantly lower, with differences of −5.12 dB, −4.24 dB, and −3.49 dB, respectively.

Topographical analyses further revealed paradigm-specific spatial distributions (Fig. 5). Text stimulation engaged bilateral ventral-stream regions near the auricular area, where neural sources are spatially closer to ear electrodes. This anatomical proximity facilitated stronger SSVEP response detection in ear-EEG despite the reduced luminance intensity of text stimuli. Importantly, the second harmonic of text responses showed a markedly different spatial distribution compared with the fundamental, with activity concentrated in the occipital cortex. This shift in activation pattern explained the rapid SNR decline at higher harmonics.

In contrast, luminance stimulation produced fundamental responses that peaked at electrode Oz and gradually declined toward the ear region, accounting for weaker ear-EEG SNR at the fundamental frequency. The second harmonic, however, displayed a broader and more diffuse spatial pattern, resulting in the highest SNR at that harmonic order.

Collectively, these results demonstrate that text and luminance paradigms evoke distinct harmonic response patterns, both in spectral distribution and spatial topography. These differences not only shape the harmonic SNR trends observed in ear-EEG versus scalp EEG, but also directly influence the classification performance across paradigms.

### Parameter Optimization

Encoding and decoding parameters played a critical role in determining BCI performance. Based on the frequency-sweeping experimental data, stimulus parameters (frequency, phase) and decoding parameters (number of sub-bands, number of electrodes, and filter bank configurations) were systematically optimized using a grid search. Classification accuracy was assessed via leave-one-out cross-validation, with 11 trials serving as the training set.

To optimize the 12-target text paradigm, we performed a grid search over (i) candidate 12-frequency sets within 4–8 Hz generated by a sliding window with a 0.2 Hz step (from 4.0–6.2 Hz up to 5.8–8.0 Hz) and (ii) phase intervals ranging from 0.05π to 2π. As shown in Fig. 7A, classification performance varied by over 10% under different frequency-phase combinations, with higher accuracy observed at lower frequencies and smaller phase differences. The highest accuracy was achieved in the 4.6–6.8 Hz range using a 0.25π phase difference, which used in12-targets paradigm.

**Fig. 7.**
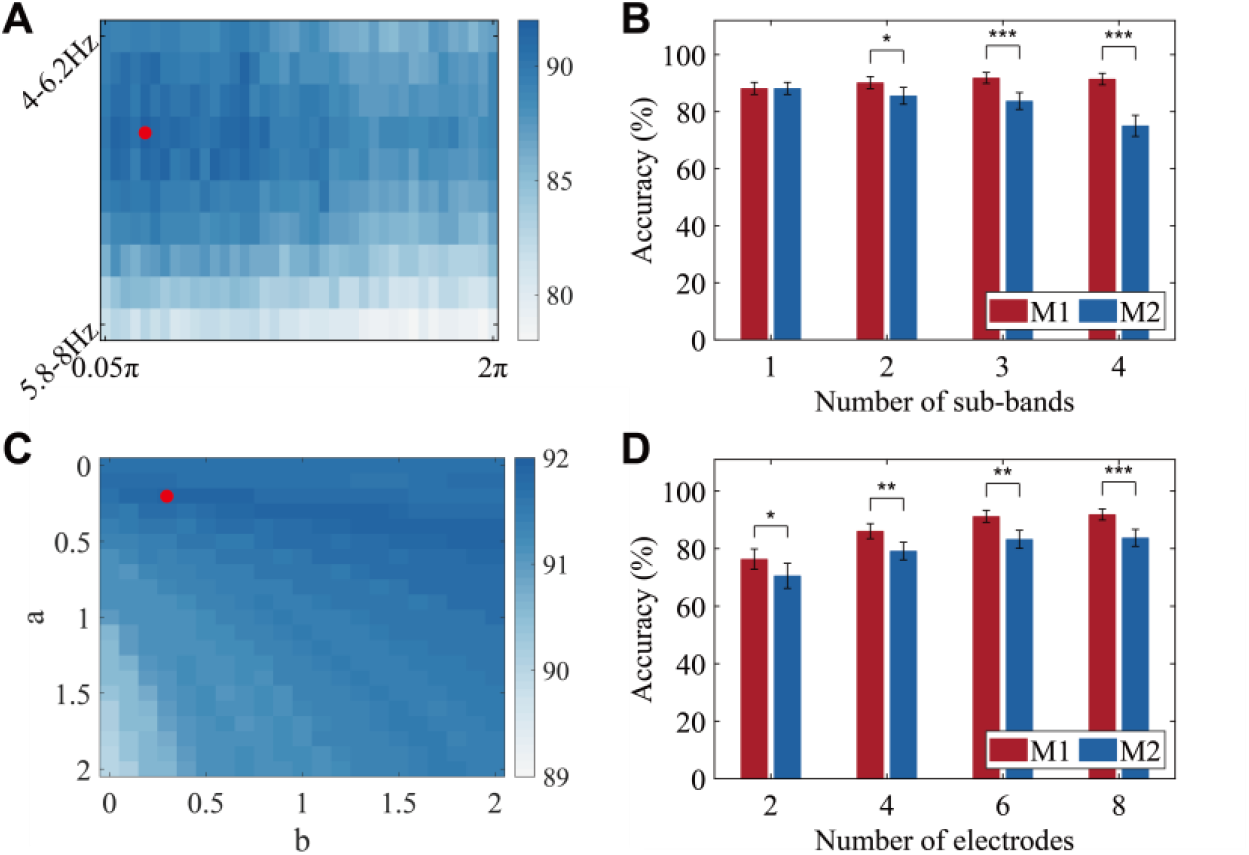
Optimization of encoding and decoding parameters under text-based stimulation. (A) Classification accuracy for a 12-target paradigm across different frequency–phase combinations. The red circle marks the combination yielding the highest accuracy. (B) Classification accuracy under different sub-band configurations. (C) Classification accuracy across varying weighting coefficients between sub -bands, with the red circle indicating the highest-performing combination.(D) Influence of electrode number on classification accuracy. M1 and M2 represent two types of sub-band configurations. (paired t-test, *p<0.05, **p<0.01, *** p<0.001).

Given the distinct spatial response patterns of text-evoked SSVEPs across harmonic frequencies, two filter-bank configurations (M1 and M2) were evaluated. As illustrated in Fig. 7B, incorporating multiple sub-bands yielded clear benefits. The M1 configuration consistently outperformed M2, achieving a peak classification accuracy of 86.4% with three sub-bands. By contrast, performance under the M2 configuration declined as the number of sub-bands increased. Moreover, as shown in Fig. 7C, further optimization of the weighting coefficients across sub-bands (a = 0.2, b = 0.3) resulted in additional gains in classification accuracy.

Additionally, as shown in Fig. 7D, we examined the influence of electrode number on classification accuracy by incrementally adding pairs of electrodes from anterior to posterior sites. As electrode count increased, classification accuracy improved markedly from 76.34%-91.85%. Increasing the number of electrodes from six to eight yielded only a slight, non-significant improvement in classification accuracy (paired *t*-test, *p* > 0.05). Notably, under every tested electrode configuration, the M1 filter bank consistently outperformed M2.

### Performance Comparison

Fig. 8 provides a detailed comparison of classification accuracy and ITR for the luminance and text paradigms over time windows. For scalp EEG, both paradigms show rapid increases in classification accuracy with shorter windows, although text-based accuracy initially lags behind luminance. By 0.8 s, the accuracies converge, and beyond 1 s both paradigms reach saturation. In terms of ITR, both paradigms exhibit an initial rise followed by a decline; luminance stimulation peaks at 0.3 s (189.48 bits/min), whereas text stimulation peaks at 0.5 s (163.66 bits/min). This lag is attributable to the lower encoding frequency of text stimuli, which encodes less information over short time windows. No participants exhibited “BCI illiteracy” under either paradigm when using scalp-EEG.

**Fig. 8.**
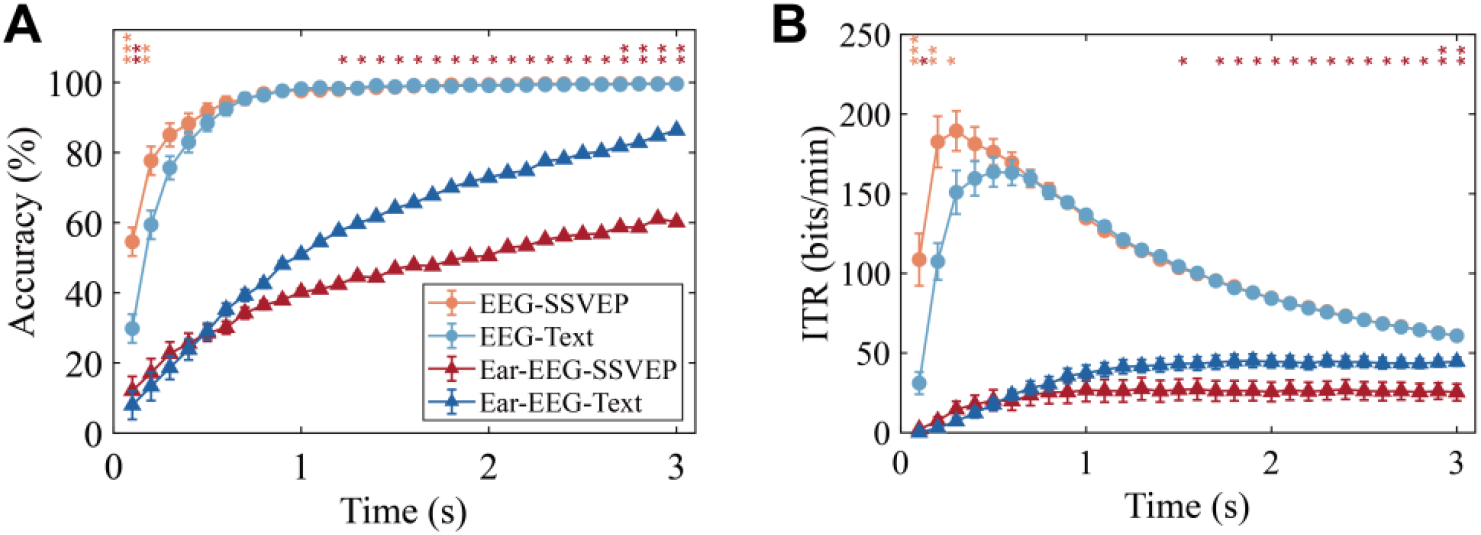
Performance of the 12-target paradigm as a function of stimulation duration. (A) Classification accuracy and (B) ITR for luminance-SSVEP and text-sequence paradigms using EEG and ear-EEG (mean ± standard error across participants). Statistical markers indicate within-subject paired t-test between the two paradigms under the same electrode type at each stimulation duration: Ear-EEG-Text vs. Ear-EEG-SSVEP (red) and EEG-Text vs. EEG-SSVEP (orange), if marked. Significance levels are *p < 0.05, **p < 0.01, ***p < 0.001.

For ear-EEG, classification accuracy improves with longer stimulation durations in both paradigms. However, text stimulation surpasses luminance after 0.5 s, showing a statistically significant accuracy advantage at 1.2 s (paired t-test, p < 0.05) and a marked ITR divergence after 1.5 s (paired t-test, p < 0.05). Luminance stimulation achieves earlier ITR saturation (0.7 s), whereas text stimulation requires 0.9 s but ultimately attains 44.59 ± 10.50 bits/min—significantly exceeding luminance-based values. This advantage corresponded to a large within-subject effect size (paired Cohen’s d = 0.86 for ITR at 3 s). Although extending the stimulation period further enhances classification accuracy in both paradigms, simply lengthening the time window does not necessarily improve the ITR. Notably, ear-EEG performance remains substantially lower than that of scalp EEG for both paradigms, indicating the need for continued optimization of ear-EEG decoding efficiency.

### Online Performance

Due to the distinct response patterns of text and luminance stimulation paradigms, considerable performance differences emerged in the online experiments. As shown in Table 1, under text stimulation with a 3 second duration, the 12-target ear-EEG BCI system achieved an average accuracy of 86.37 ± 9.61% and an ITR of 44.59 ± 10.50 bits/min. The top-performing participant reached 100% accuracy and ITR over 60 bits/min. Even the participant with the lowest performance attained 69.17% accuracy and 27.89 bits/min—surpassing the average under luminance stimulation. By contrast, under luminance stimulation, participant performance varied widely: the poorest accuracy was only marginally above chance, corresponding to an ITR of merely 2.81 bits/min, whereas the best participant achieved 98.33% accuracy.

**TABLE I.** Online Performance.

Online Performance
| Subject | Text |  | Luminance |  |
| --- | --- | --- | --- | --- |
|  | Accuracy (%) | ITR (bits/min) | Accuracy (%) | ITR (bits/min) |
| Sub1 | 100 | 61.46 | 40.83 | 9.64 |
| Sub2 | 88.33 | 45.63 | 72.50 | 30.60 |
| Sub3 | 82.50 | 39.61 | 25.00 | 3.07 |
| Sub4 | 95.83 | 54.70 | 98.33 | 58.37 |
| Sub5 | 82.50 | 39.61 | 80.83 | 38.01 |
| Sub6 | 73.33 | 31.30 | 85.00 | 42.11 |
| Sub7 | 99.17 | 59.77 | 24.17 | 2.81 |
| Sub8 | 96.67 | 55.87 | 88.33 | 45.63 |
| Sub9 | 88.33 | 45.63 | 50.00 | 14.66 |
| Sub10 | 91.67 | 49.42 | 39.17 | 8.82 |
| Sub11 | 82.50 | 39.61 | 82.50 | 39.61 |
| Sub12 | 69.17 | 27.89 | 37.50 | 8.03 |
| Sub13 | 76.67 | 34.18 | 26.67 | 3.62 |
| Sub14 | 82.50 | 39.61 | 91.67 | 49.42 |
| Mean | 86.37 | 44.59 | 60.18 | 25.31 |
| ±Std | ±9.61 | ±10.50 | ±27.77 | ±19.93 |

Compared with luminance modulation, text-sequence stimulation increased the ITR by 76.18% (paired t-test, p = 0.0068) and markedly reduced the proportion of low-performing users in the ear-EEG BCI. Using 70% accuracy as a practical usability threshold, all participants achieved near or above 70% accuracy with text stimuli, whereas 50% of participants remained below 70% under luminance modulation. In addition, participant Sub13 (57 years old, with no prior BCI/EEG experience) achieved an ITR of 34.18 bits/min with text stimulation but performed poorly under luminance stimulation; this case is reported only as a preliminary feasibility observation rather than evidence of age-independence.

To assess the robustness of the group-level conclusions, we repeated the analysis after excluding Sub13. The results remained consistent: Text achieved 87.12 ± 9.57% accuracy and 45.39 ± 10.47 bits/min ITR, whereas luminance achieved 62.76 ± 27.10% accuracy and 26.98 ± 19.70 bits/min ITR, corresponding to a 68.23% ITR increase. The paired comparisons remained significant (p = 0.014 for ITR; p = 0.0117 for accuracy, paired t-test). Together, these results suggest that the proposed text-sequence paradigm may provide more robust online spelling performance than conventional luminance modulation. However, because only one older participant was included, further studies with a broader age distribution are needed to draw definitive conclusions regarding age effects.

### Experience Evaluation

In the text sequence stimulation paradigm, the effective stimulation area was limited (as defined by the red box boundary in Fig. 2C), the average area of text stimuli occupied only about 14% of the luminance stimulation. Specifically, the luminance of individual characters ranged from 16.5 to 143.8 cd/m^2^, with an average luminance of 100.6 cd/m^2^. In contrast, the luminance-modulated SSVEP paradigm exhibited a much broader dynamic range, varying from 0 to 423 cd/m^2^. Such a substantially larger stimulation area and higher luminance variation are more likely to induce visual fatigue, which in turn can further degrade BCI performance, leading to a negative feedback loop.

According to the user experience ratings shown in Fig. 9, the luminance-modulated SSVEP paradigm scored below 2, whereas the text-based sequence paradigm scored around 4. This outcome indicates that, relative to luminance modulation, the text sequence stimulation method offers enhanced comfort, reduced flicker perception, and greater user preference. The questionnaire was intended as a within-subject comparative assessment of visual experience rather than a validated clinical scale.

**Fig. 9.**
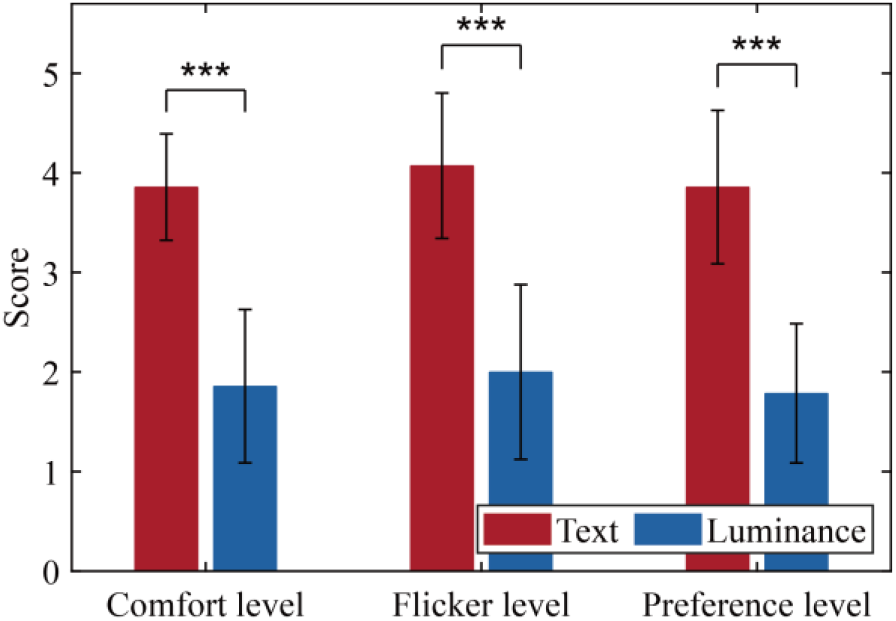
Experience evaluation of text and luminance paradigms. Statistical significance levels are indicated as ***p<0.001. Error bar represents the standard error.

## 4 Discussion

This study demonstrates that text sequence stimulation significantly enhances both performance and user ergonomics in ear-EEG-based BCIs. The proposed paradigm achieved an ITR of44.59 ± 10.50 bits/min, surpassing conventional luminance modulation by76.18%. This improvement stems from two key innovations: (1)text-specific neural activation patternsoptimized for ear-EEG’s spectral characteristics (4.6–6.8 Hz, 0.25π phase interval), and (2)filter banks (M1 configuration)that exploit response differences across frequency bands. Notably, text stimulation eliminated BCI illiteracy, with all participants achieving near or above 70% accuracy—a critical advancement for applicability.

As summarized in Table 2, most existing ear-EEG SSVEP studies rely on low-frequency luminance-modulated stimuli. Although they achieve relatively high ITRs, these stimuli can induce visual fatigue and offer limited spatial alignment with ear electrodes[4,7]. While alternative paradigms, such as color modulation and bilateral phase coding, may improve comfort or boost SNRs, they often compromise either target capacity or ITR[12,18]. For instance, depth-modulated stimuli achieved 91% accuracy but were limited to only two targets[19]. By contrast, the text paradigm leverages ventral pathway activation near the ear, combining reduced flicker intensity (14% of luminance stimuli area) with efficient multi-target encoding.

**TABLE II.**
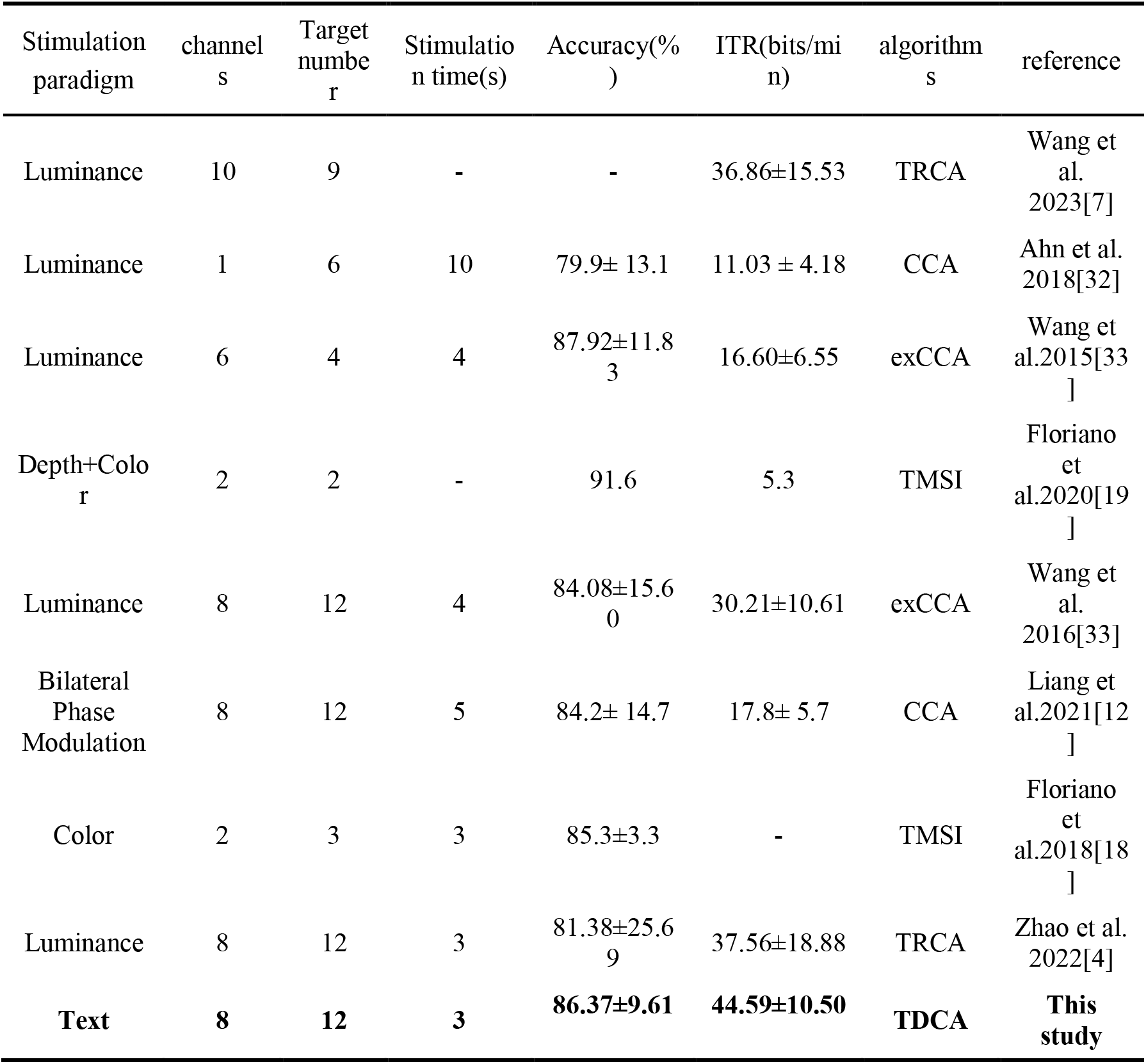
Previous Studies to ear-EEG based SSVEP BCI.

| Stimulation paradigm | channels | Target number | Stimulation time(s) | Accuracy(%) | ITR(bits/min) | algorithms | reference |
| --- | --- | --- | --- | --- | --- | --- | --- |
| Luminance | 10 | 9 | - | - | 36.86±15.53 | TRCA | Wang et al. 2023[7] |
| Luminance | 1 | 6 | 10 | 79.9± 13.1 | 11.03 ± 4.18 | CCA | Ahn et al. 2018[32] |
| Luminance | 6 | 4 | 4 | 87.92±11.83 | 16.60±6.55 | exCCA | Wang et al.2015[33] |
| Depth+Color | 2 | 2 | - | 91.6 | 5.3 | TMSI | Floriano et al.2020[19] |
| Luminance | 8 | 12 | 4 | 84.08±15.60 | 30.21±10.61 | exCCA | Wang et al. 2016[33] |
| Bilateral Phase Modulation | 8 | 12 | 5 | 84.2± 14.7 | 17.8± 5.7 | CCA | Liang et al.2021[12] |
| Color | 2 | 3 | 3 | 85.3±3.3 | - | TMSI | Floriano et al.2018[18] |
| Luminance | 8 | 12 | 3 | 81.38±25.69 | 37.56±18.88 | TRCA | Zhao et al. 2022[4] |
| **Text** | **8** | **12** | **3** | **86.37±9.61** | **44.59±10.50** | **TDCA** | **This study** |

BCI illiteracy—a major barrier to widespread adoption—is particularly pronounced in ear-EEG systems using occipital-dominant luminance stimuli[12,16], where approximately 50% of users fail to achieve 70% accuracy. In our study, all participants exhibited robust decoding performance with scalp EEG. However, performance varied substantially across individuals in ear-EEG systems. Previous ear-EEG studies often lack detailed reports of single-subject performance. In bilateral phase-modulation paradigms, 5 out of 15 participants failed to reach 70% accuracy, even though optimization improved the overall average[12].

These individual differences suggest that ear-EEG BCI illiteracy may largely arise from spatial misalignment between neural sources and recording sites. Text stimuli shift neural responses closer to the ear electrodes, enabling all participants to reach or exceed the 70% accuracy threshold. The elimination of BCI illiteracy in ear-EEG demonstrates the potential of text sequence paradigms for inclusive deployment in assistive communication and environmental control applications[14,18].

From a rehabilitation perspective, these findings suggest that paradigm-level stimulus design may play a critical role in improving the usability of ear-EEG BCIs in populations with severe motor impairment. Future studies will evaluate the proposed paradigm in target user groups and focus on practical metrics such as setup time, visual fatigue, prolonged use stability, and caregiver-assisted operation. The present results in healthy participants therefore represent a necessary step toward rehabilitation-oriented validation rather than a final clinical endpoint.

In recent years, SSVEP decoding algorithms have made substantial progress. Our understanding of EEG signals in the frequency domain has evolved from canonical correlation analysis (CCA) using sine/cosine templates[34], to filter bank CCA (FBCCA)[26], and finally to spatiotemporal equalization methods that optimize both the signal and noise subspaces[35]. Each advancement has consistently enhanced system performance. However, ear-EEG SSVEP responses differ markedly from those of scalp EEG in both spatial and frequency domains, which impacts decoding effectiveness. Relatively few studies have specifically optimized algorithms for ear-EEG. In this work, we redesigned the filter bank configuration (M1), thereby improving overall system performance.

Despite these advances, ear-EEG ITR (44.59 bits/min) still falls short of that achieved by scalp EEG-based systems. Three strategies could help bridge this gap. First, increased electrode density: Current ear-EEG setups use only eight electrodes, limiting spatial resolution. Flexible electronics could support dense, conformal electrode arrays to enhance signal fidelity[7,36]. Second, improved encoding methods: Future research will explore semantic content and character-specific neural responses to refine stimulus personalization[37,38]. Third, robust decoding algorithms: Proximity to facial muscles increases susceptibility to EMG noise, and neural networks could more effectively separate SSVEP components from background activity[39,40]. Through these enhancements, text-based ear-EEG BCIs can continue to evolve, bringing us closer to high-performance, user-friendly systems suitable for widespread, real-world applications.

This study has several limitations. The cohort size is moderate (offline: n=15; online: n=14) and does not establish population-level generalizability. In addition, all experiments were conducted in a controlled laboratory setting, and robustness in naturalistic environments (e.g., motion, ambient-light variability, and daily-life distractions) was not evaluated. Future work will validate the paradigm in larger and more diverse cohorts and conduct field-oriented tests to assess real-world robustness and usability (e.g., longer use, comfort/fatigue, setup time, and electrode stability).

## 5 Conclusions

This study demonstrates that text-sequence stimulation can substantially enhance ear-EEG BCI performance, achieving an average ITR of 44.59 10.50 bits/min, compared with only 25.31±19.93 bits/min under luminance stimulation, corresponding to a 76.18% improvement. At the same time, the text paradigm markedly reduced the required stimulation area, thereby improving user ergonomics. Critically, text stimuli eliminated the “BCI illiteracy” problem, with all participants achieving accuracy levels at or above 70%. These findings validate text-based stimulation as a practical and efficient paradigm for wearable ear-EEG systems, striking a balance between high performance and user-friendliness.

## 6 List of abbreviations

(BCIs): Brain–computer interfaces
(EEG): electroencephalography
(SSVEPs): steady-state visual evoked potentials
(ITRs): information transfer rates
(SNRs): signal-to-noise ratios
(TDCA): task-discriminant component analysis

## 7 Author contributions

Conceptualization: Xiaoyang Li.

Investigation: Xiaoyang Li, Zhuoran Xu.

Methodology: Xiaoyang Li, Bowen Li.

Writing – Original Draft: Xiaoyang Li, Zhuoran Xu.

Review & Editing: Yijun Wang, Xiaorong Gao.

Visualization: Xiaoyang Li, Zhuoran Xu, Bowen Li.

Supervision: Yijun Wang, Xiaorong Gao.

Funding Acquisition: Yijun Wang, Xiaorong Gao.

All authors read and approved the final manuscript.

## 8 Acknowledgment

The authors would like to thank Changxing Huang, Yonghao Song and Yuzhen Chen from Tsinghua University for the helpful discussion.

## 9 Ethics approval and consent to participate

The study was approved by the Tsinghua University Ethics Committee (No: THU01-20240030), and informed consent was obtained.

## 11 Consent for publication

Not applicable.

## 13 Availability of data and materials

Data will be made available on request.

